# Testing the ‘caves as islands’ model in two cave-obligate invertebrates with a genomic approach

**DOI:** 10.1101/2020.04.08.032789

**Authors:** Andras Balogh, Lam Ngo, Kirk S. Zigler, Groves Dixon

## Abstract

Caves offer selective pressures that are distinct from the surface. Organisms that have evolved to exist under these pressures typically exhibit a suite of convergent characteristics, including a loss or reduction of eyes and pigmentation. As a result, cave-obligate taxa, termed troglobionts, are no longer viable on the surface. This circumstance has led to a “caves as islands” model of troglobiont evolution that predicts extreme genetic divergence between cave populations even across relatively small areas. An effective test of this model would involve (1) common troglobionts from (2) nearby caves in a cave-dense region, (3) good sample sizes per cave, (4) multiple taxa, and (5) genome-wide characterization. With these criteria in mind, we used RAD-seq to genotype an average of ten individuals of the troglobiotic spider *Nesticus barri* and the troglobiotic beetle *Ptomaphagus hatchi*, each from four closely located caves (ranging from 3-13 km apart) in the cave-rich southern Cumberland Plateau of Tennessee, USA. Consistent with the caves as islands model, we find that populations from separate caves are indeed highly genetically isolated. In addition, nucleotide diversity was correlated to cave length, suggesting that cave size is a dominant force shaping troglobiont population size and genetic diversity. Our results support the idea of caves as natural laboratories for the study of parallel evolutionary processes.

## Introduction

Caves are unique habitats with environmental conditions fundamentally distinct from the surface. The most conspicuous of these is the complete absence of light, precluding the use of visual cues for hunting, foraging, locating mates, and evading predators (Rétaux and Casane 2013). Moreover, as photosynthesis is not possible, cave communities depend nearly entirely on trophic input from the surface. Caves are also typically more stable in temperature and humidity than surface habitats (Culver and Pipan 2019). As a result, caves offer opportunities for extensive evolutionary change (Poulson and White 1969).

Some organisms have evolved under these conditions to the extent that they are never found outside of caves. These organisms, termed troglobionts (Culver and Pipan 2019), often bear a suite of distinctive characteristics, including loss or reduction of eyes, pigment loss, elongated appendages, improved non-visual sensory mechanisms, reduced metabolic rates, longer lifespans, and lower rates of reproduction (Poulson and White 1969; Peck 1986; Culver and Pipan 2019). Many of these phenotypes have obvious tradeoffs for fitness on the surface. For instance, pigment loss, selectively neutral or possibly even advantageous in the cave environment (Polo-Cavia and Gomez-Mestre 2017), will decrease crypsis on the surface, a trait known to undergo particularly strong purifying selection (Cook and Saccheri 2013). Intolerance to variation in temperature and humidity may also preclude surface viability (Culver and Pipan 2019). Hence, the surface is a hostile environment to troglobionts. With this idea in mind, Culver and Pipan (2019) pointed out that caves are like islands in a sea of surface habitat.

Previous studies have shown that troglobiont migration is indeed highly limited. For instance, at the species level, endemism is less the exception than the rule (Culver et al. 2000; Niemiller and Zigler 2013). This is especially true in the eastern United States, where up to 45% of troglobionts are single- cave endemics (Christman et al. 2005). These exceptional rates of endemism are consistent with restricted gene flow and frequent speciation. Population genetic studies lend further support. Examining COI in the troglobiotic spider *Nesticus barri*, Snowman et al. (2010) found extensive haplotype divergence and limited sharing of haplotypes between caves, indicating that migration was minimal to nonexistent over distances greater than 15 km. Another study, examining COI in several troglobionts, including *N. barri* and the beetle *Ptomaphagus hatchi*, provided similar findings (Dixon and Zigler 2011), indicating that restricted migration is likely general to terrestrial troglobionts. In contrast, some aquatic troglobionts show high connectivity between caves, and unexpectedly large population sizes (Buhay and Crandall 2005; Buhay et al. 2007). The contrast between aquatic and terrestrial troglobionts may reflect broader aquatic subterranean connectivity than terrestrial (Porter 2007). Hence, for small terrestrial troglobionts caves indeed often resemble islands.

Here, we sought to critically evaluate the caves as islands hypothesis for two cave-obligate invertebrates. To emphasize the severity of isolation imposed by troglobiont ecology and life history, we sampled individuals from caves located closely together in the cave-dense region of the southern Cumberland Plateau, one of the most biodiverse karst areas in the United States (Culver et al. 2000; Christman and Culver 2001; Niemiller and Zigler 2013) (Fig. 1; Table 1). To ensure the generality of the hypothesis, we focused on two species with distinct natural histories: the spider *Nesticus barri* and the beetle *Ptomaphagus hatchi. Nesticus barri* is part of a complex of 28 species found across the southeastern United States (Hedin 1997). Their tendency to live in dark, moist habitats has led to numerous instances of cave habitation, with roughly one third of species in the group either troglophiles (frequent cave dwellers that are also found on the surface) or troglobionts (Hedin and Dellinger 2005). *Nesticus barri* demonstrates typical troglomorphic features, lacking eyes and with reduced pigment, although it still possesses reproductive seasonality (Carver et al. 2016). The genus *Ptomaphagus* includes about 60 species in North America, again with roughly one third either troglophiles or troglobionts (Peck 1986). Diversification of the genus throughout the southern Cumberland Plateau is thought to have occurred through progressive vicariance, as the Cumberland Plateau eroded over the last six million years (Leray et al. 2019). Like other troglobiotic *Ptomaphagus, P. hatchi* has greatly reduced eyes and is wingless. The southern Cumberland Plateau is one of the most cave-rich regions in North America, with more than 4000 caves known from a six-county area in southern Tennessee and northeast Alabama (Zigler et al. 2014). On the southern Cumberland Plateau, *N. barri* and *P. hatchi* have largely overlapping ranges, with each species known from dozens of caves (Snowman et al. 2010; Leray et al. 2019), and both species are common in the caves they inhabit. Finally, where previous studies made use of one, or at most a handful loci, we take a genome wide approach using 2bRAD (Wang et al. 2012). This method has the advantage of interrogating thousands of loci across the genome, allowing for more confident estimates of population divergence and neutral diversity (Rokas and Abbot 2009; Nunziata and Weisrock 2018). This is the first study to investigate these species at the genomic scale.

**Table 1:** Cave information

| Cave | Abbreviation | Survey # | Length (m) | Watershed | Sample size |  |
| --- | --- | --- | --- | --- | --- | --- |
|  |  |  |  |  | *N. barri* | *P. hatchi* |
| Solomon's Temple Cave | ST | TFR26 | 370 | Upper Elk River | 12 | 6 |
| Sewanee Blowhole | SB | TFR91 | 1219 | Upper Elk River | 8 | 7 |
| Grapevine Cave | GV | TFR423 | 490 | Guntersville Lake | 10 | 12 |
| Buggytop Cave | BT | TFR16 | 3142 | Guntersville Lake | 16 | 7 |

**Figure 1:**
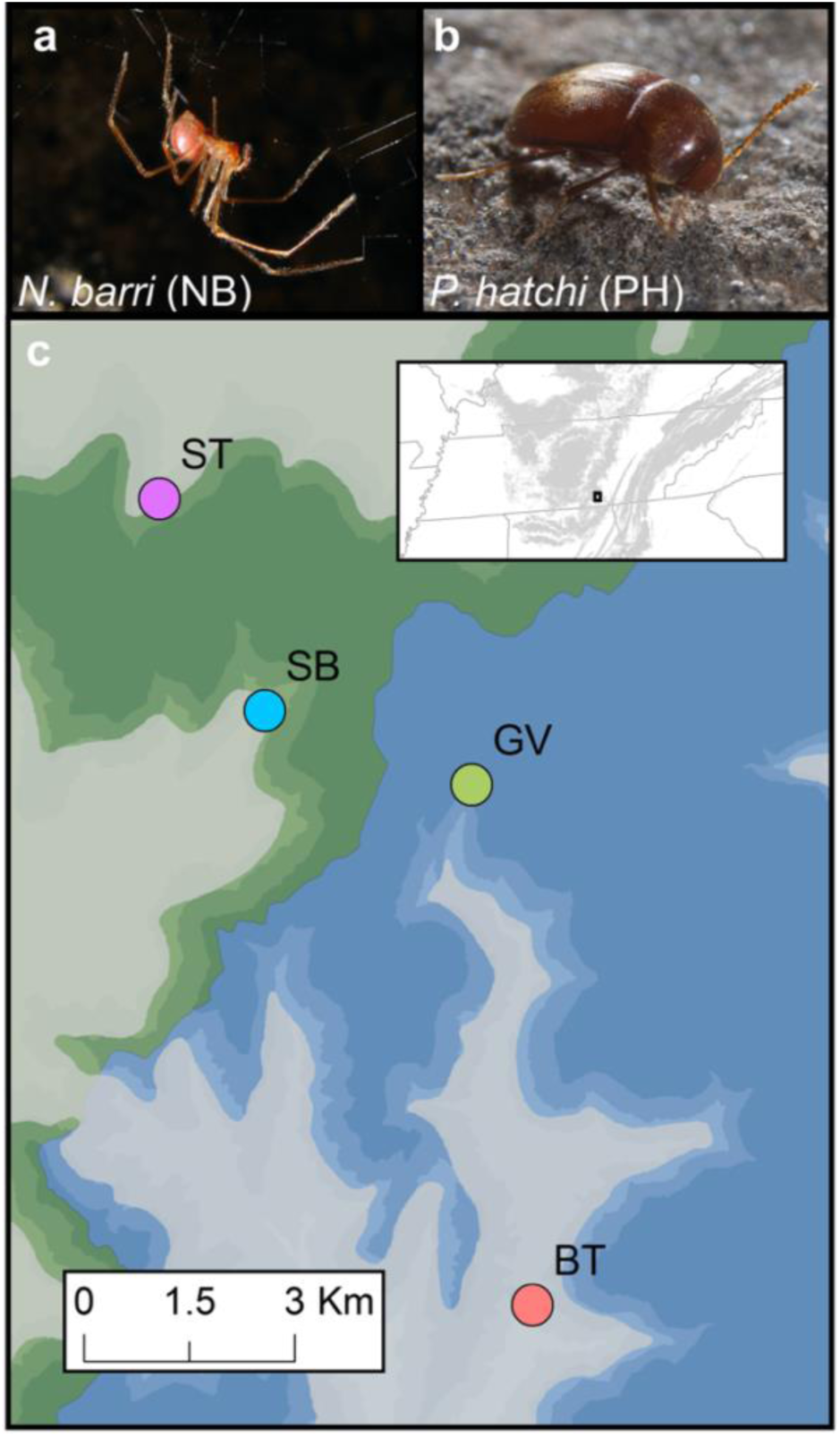
Study location and abbreviations. (a) *Nesticus barri* (NB) (b) *Ptomaphagus hatchi* (PH) (c) Map of sample cave locations: Buggytop (BT); Grapevine (GV); Sewanee Blowhole (SB); Solomon’s Temple (ST). The Upper Elk River watershed is indicated by green coloration and the Guntersville Lake watershed is indicated by blue coloration. Shading intensity indicates elevation with higher elevations in darker tones. Photos by (a) Alan Cressler and (b) Michael Slay.

## Materials and methods

### Sampling

We collected specimens from four caves on the edge of the southern Cumberland Plateau in Franklin County, Tennessee (Table 1). The caves were chosen based on proximity, location, and previous knowledge of the presence of *Nesticus barri* and *Ptomaphagus hatchi* (Dixon and Zigler, 2011; Wakefield and Zigler, 2012). Distances between the caves ranged from 3-13 km (Fig. 1). The caves are distributed across two adjacent watersheds, with Solomon’s Temple (ST) and Sewanee Blowhole (SB) in the Upper Elk River watershed that drains to the north and west of the study area, whereas Grapevine Cave (GV) and Buggytop Cave (BT) are in the Guntersville Lake watershed that drains the study area to the south (Fig. 1; Table 1).

Sampling was conducted between 21 September and 25 October 2018. We collected *Ptomaphagus* and *Nesticus* by hand during visual encounter surveys of the caves. An initial survey of Buggytop Cave yielded only a few *Ptomaphagus*, so food baits (tuna) were placed in the cave for 24 hours and live specimens were subsequently collected at the baits. Sample size per cave ranged from 6-16 individuals per species (Table 1). Specimens were placed into 100% EtOH in the field and subsequently stored at - 20°C. Sampling was permitted by the Tennessee Wildlife Resources Agency (Permit #1385) and the Tennessee Department of Environment and Conservation (Permit #2013-026).

### Library preparation

Most of the DNA extractions were performed using the entire body of the individual. If a particular sample seemed large enough (mainly applied to *Nesticus barri*) the legs were saved while the cephalothorax and abdomen were used. QIAGEN’s DNeasy Blood & Tissue Kit (cat. no. 69504 or 69506) was used following the kit’s protocol with the exception of using 50 µl Buffer AE for elution. Concentrations of each DNA isolation were initially checked by nanodrop and confirmed with the Quant-IT Picogreen DS DNA assay (Life Technologies cat. no. P7589). The 2b-RAD library preparation was carried out as described previously (Wang et al. 2012; Dixon et al. 2015; Matz et al. 2018; Matz 2019). Briefly, DNA isolations were normalized to ∼12.5 ng/µl. Samples with concentrations lower than 12.5 ng/µl DNA (∼30 samples), were fully dehydrated in a vacuum centrifuge and resuspended to a target concentration of ∼12.5 ng/µl. Digestion reactions had concentrations of 1x NEB buffer #3 and 10 µM SAM mixed with 1 total U of BcgI restriction enzyme and 50 ng genomic DNA in a total volume of 6 µl. Digests were incubated at 37°C for 1 hour followed by 20 minutes at 65°C for heat inactivation. Ligation reactions had concentrations of 1x T4 ligase buffer, and 0.25 µM each adapter with 400 total U of T4 DNA ligase and 6 µl of digested DNA in a total volume of 20 µl. Ligation reactions were incubated at 4°C overnight, followed by 20 minutes at 65°C for heat inactivation. Inclusion of internal barcodes in the i7 adapter allowed for pooling sets of samples at this point. Amplification and additional barcoding reactions were performed on these pools. These reactions had concentrations of 312 µM each dNTP, 0.2 µM each of p5 and p7, 0.15 µM appropriate TruSeq-Un primer, 0.15 µM appropriate primer, 1x Titanium Taq buffer and 1x Taq polymerase mixed with 4 µl of pooled ligation in a total volume of 20 µl. Thermocycler conditions were 70°C for 30 seconds, followed by 14 cycles of 95°C for 20 seconds, 65°C for 3 minutes, and 72°C for 30 seconds. The final library was pooled into a single tube and sequenced at the University of Texas at Austin’s Genome Sequencing and Analysis Facility on a single lane on the Illumina Hiseq 2500 platform.

### Data processing

Raw reads were trimmed and demultiplexed based on internal ligation barcodes using custom perl scripts (Matz 2019). Reads were quality filtered using fastq_quality_filter from the Fastx Toolkit (http://hannonlab.cshl.edu/fastx_toolkit/). Generation of de novo loci was performed using cd-hit (Li and Godzik 2006) and custom perl scripts as described previously (Wang et al. 2012; Dixon et al. 2015; Matz et al. 2018; Matz 2019). These tags were assembled into a reference genome with 30 equally sized pseudo-chromosomes for mapping. Re-mapping of reads to these de novo loci was done with bowtie2 (Langmead and Salzberg 2012). Sorting and indexing of bam files in preparation for genotyping was done with samtools (Li et al. 2009).

### Genotype analyses

We analyzed the genotype data in two ways. The first method depended on hard genotype calls, in which the genotype of each individual at each site is either called exactly, or filtered to missing data based on arbitrary cutoffs. While simpler to implement, these hard genotype calls can introduce biases, because they can fail to capture statistical uncertainty inherent to individual genotypes from New-Generation Sequencing (NGS) data (Nielsen et al. 2012). A second, alternative approach is to estimate sample allele frequency spectra directly from base calls and quality metrics in the alignment data, allowing for population genetic inferences without making individual genotype calls (Nielsen et al. 2012). We processed hard genotype calls primarily using VCFtools (Danecek et al. 2011). Estimation of genotype likelihoods, allele frequency spectra, and additional population genetic inferences were implemented using Angsd (Korneliussen et al. 2014). Throughout the manuscript we emphasize the results produced using Angsd, using analyses from hard genotype calls primarily for corroboration. All steps used for both sets of analyses, along with scripts for statistical analysis and figure generation are included in the git repository (Dixon 2019).

Hard genotype calls were made as described previously (Dixon et al. 2019) using mpileup and bcftools (Li 2011). Genotype calls with a depth lower than 2, as well as indels, singletons, sites with more than two alleles or less than 75% of samples genotyped were removed using VCFtools (Danecek et al. 2011). Sites with excess heterozygosity (p-value less than 0.1) (likely paralogs) were removed based on the --hardy output from VCFtools.

For the analyses performed with Angsd, quality filtering included a minimum mapping score of 20 (-minMapQ 20), a minimum quality score of 32 (-minQ 32), and minimum representation among samples > 80% (-minInd). Filters based on p-values, including the strand bias p-value (-sb_pvalue)(Guo et al. 2012), the hetbias p-value (-hetbias_pval)(Vieira et al. 2013), and SNP pvalue (-snp_pval), were set to 0.05.

### Summarizing genetic variation

To summarize genetic variation, we used Angsd to calculate pairwise differences between samples using the -IBS 1 option and a minimum minor allele frequency of 1% (-minMaf). Here pairwise distance between samples *i* and *j* (*d*_*ij*_) is calculated as:

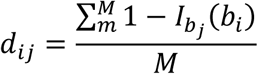

Where *M* is the total number of sites with at least 1 read from each individual, and *1 - I*_*bj*_*(b*_*i*_) is the indicator function which is equal to one when the two individuals have the same base and zero otherwise (Korneliussen 2013). This distance matrix was used for hierarchical clustering and multidimensional scaling. Admixture analysis was performed on genotype likelihoods output by Angsd using NGSadmix (Skotte et al. 2013).

### Genetic differentiation between populations based on SNP calls

We estimated F_ST_ for each pair of populations using Angsd (Fumagalli et al. 2013; Korneliussen et al. 2014). This method computes the posterior expectation of genetic variance between populations (designated A), and total expected variance (designated B). These values (A and B) are closely related to the alpha and beta estimates described in Reynolds et al. (1983). The unweighted F_ST_ is computed as the mean of the per-site ratios of A and B and the weighted FST is computed as the ratio of the sum of As to the sum of Bs (Korneliussen 2013). The unweighted and weighted F_ST_ values reported in Table 2 and Figure 4 are these estimates computed from all sites in the dataset. Based on communications that unweighted F_ST_ values tend to be too noisy (ANGSD 2014) we focus on the weighted F_ST_ values.

**Table 2:** Estimates of pairwise genetic differentiation between caves. ‘Weighted’ and ‘unweighted’ indicate the F_ST_ estimates computed with Angsd. ‘Weir’ indicates Weir and Cockerham’s F_ST_ estimate computed with VCFtools. ‘d_XY_’ indicates pairwise genetic distance computed from VCFtools allele frequencies using the equation given in the methods section.

| pair | watershed | species | weighted | unweighted | Weir | $d_{xy}$ |
| --- | --- | --- | --- | --- | --- | --- |
| SB-ST | same | *N. barri* | 0.33 | 0.12 | 0.20 | 0.14 |
| BT-GV | same | *N. barri* | 0.42 | 0.13 | 0.29 | 0.25 |
| BT-ST | different | *N. barri* | 0.42 | 0.14 | 0.33 | 0.31 |
| BT-SB | different | *N. barri* | 0.43 | 0.14 | 0.32 | 0.30 |
| GV-ST | different | *N. barri* | 0.49 | 0.15 | 0.45 | 0.33 |
| GV-SB | different | *N. barri* | 0.52 | 0.18 | 0.45 | 0.32 |
| SB-ST | same | *P. hatchi* | 0.34 | 0.18 | 0.25 | 0.21 |
| BT-GV | same | *P. hatchi* | 0.30 | 0.17 | 0.16 | 0.22 |
| BT-ST | different | *P. hatchi* | 0.34 | 0.17 | 0.27 | 0.25 |
| BT-SB | different | *P. hatchi* | 0.32 | 0.17 | 0.15 | 0.24 |
| GV-ST | different | *P. hatchi* | 0.36 | 0.19 | 0.31 | 0.25 |
| GV-SB | different | *P. hatchi* | 0.36 | 0.20 | 0.22 | 0.24 |

Pairwise F_ST_ and d_XY_ were also calculated for each pair of caves based on hard genotypes. We used VCFtools to calculate pairwise F_ST_ (Weir and Cockerham 1984), and a custom R scrip to calculate d_XY_ for unphased data as:

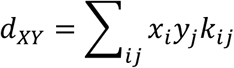

Where *x*_*i*_ and *y*_*j*_ are the frequencies of the *i*th allele from population X and the *j*th allele from population Y respectively, and *k*_*ij*_ is 1 when *i* and *j* differ, and 0 if they are the same (Hahn 2018). The Weir and Cockerham’s F_ST_ and the d_XY_ values reported in Table 2 and Fig. 4 are the averages of these statistics across all hard genotyped SNPs.

### Nucleotide diversity

To calculate nucleotide diversity using genotype likelihoods, we first generated genotype likelihoods as described above. We then used the custom python script HetMajorityProb.py (Matz et al. 2018; Matz 2019) to remove sites where the heterozygosity rate appeared higher than 50%, as these were likely paralogs spuriously lumped as single loci. We then estimated nucleotide diversity from the folded site frequency spectra using Angsd (Nielsen et al. 2012; Korneliussen et al. 2013, 2014). The value was then averaged across the 30 pseudochromsomes from the reference created during de novo locus generation described above. These averages are the values reported in Figure 5.

We also calculated nucleotide diversity (π) from hard genotype calls. Here we first determined the allele frequencies for each species in each cave using VCFtools. We then calculated expected heterozygosity (*h*) for each site as:

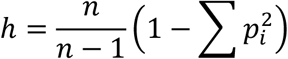

Where *n* is the number of sequences, and *p*_*i*_ is the frequency of the *i*th allele at the site. We then calculated π as the sum of expected heterozygosities across sites:

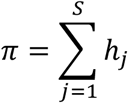

Where *S* is the number of segregating sites and *h*_*j*_ is the expected heterozygosity of the site (Hahn 2018). We report this value per site by dividing by the total number of interrogated positions. Effective population size was estimated by dividing this per site value by four times the number of mutations per site per generation (4µ). The mutation rate estimate was based on *Drosophila* (Keightley et al. 2014).

## Results

### Sequencing

Sequencing produced 236 million raw reads. After demultiplexing and PCR duplicate removal, a total of 65.8 million reads remained. Mean fold coverage per sample tended to be higher for *P. hatchi* (1.2 ± se 0.08 million reads) than *N. barri* (0.462 ± se 0.04 million reads). De novo locus generation produced 130,061 loci for *N. barri* and 264,486 loci for *P. hatchi*. Filtering of hard genotype calls from mpileup produced 7,620 biallelic SNPs for *N. barri* and 12,879 for *P. hatchi*. Filtering of genotype likelihoods produced with Angsd identified 13,629 SNPs for *N. barri* and 31,000 for *P. hatchi*. Final sample sizes for each species are given in Table 1.

### Delineating populations

For both species, the four caves harbored genetically distinct populations. Hierarchical clustering based on pairwise differences separated individuals by cave (Fig. 2a,d). Overall topology of clustering was the same for both species, with caves from the same watershed clustering together (Fig. 1; Fig. 2a,d). Multidimensional scaling produced similar results, with clear clustering by cave along the first three axes for *N. barri* (Fig. 2b,c) and the first two for *P. hatchi* (Fig. 2e,f). Both analyses indicated that SB and ST caves were more similar for *N. barri* than *P. hatchi*.

**Figure 2:**
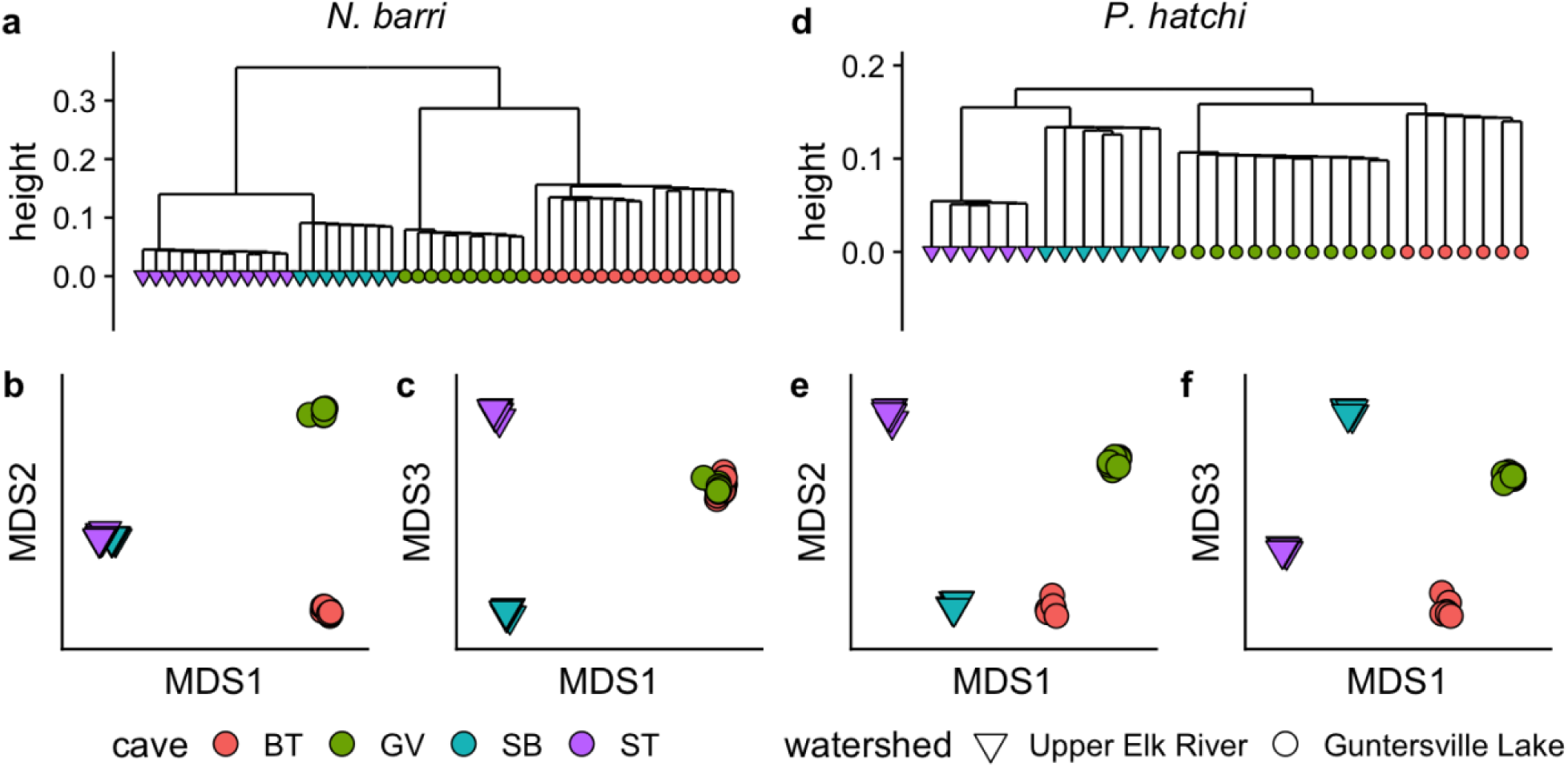
Sample clustering by cave. (a) Hierarchical clustering of *N. barri* samples based on pairwise distance. (b-c) Multidimensional scaling plots based on pairwise distance for *N. barri*. (d-f) Same plots for *P. hatchi*.

Admixture analysis further supported the caves as independent populations. For both species, when the number of ancestral populations (k) was set to four, ancestry estimates matched fully with source cave (Fig. 3). For both species, division of ancestry among five ancestral populations lead to a split of Buggytop (the largest cave) into two roughly equally sized groups. Together, these analyses indicate that the four caves are indeed distinct populations, harboring genetically distinct individuals for both species.

**Figure 3:**
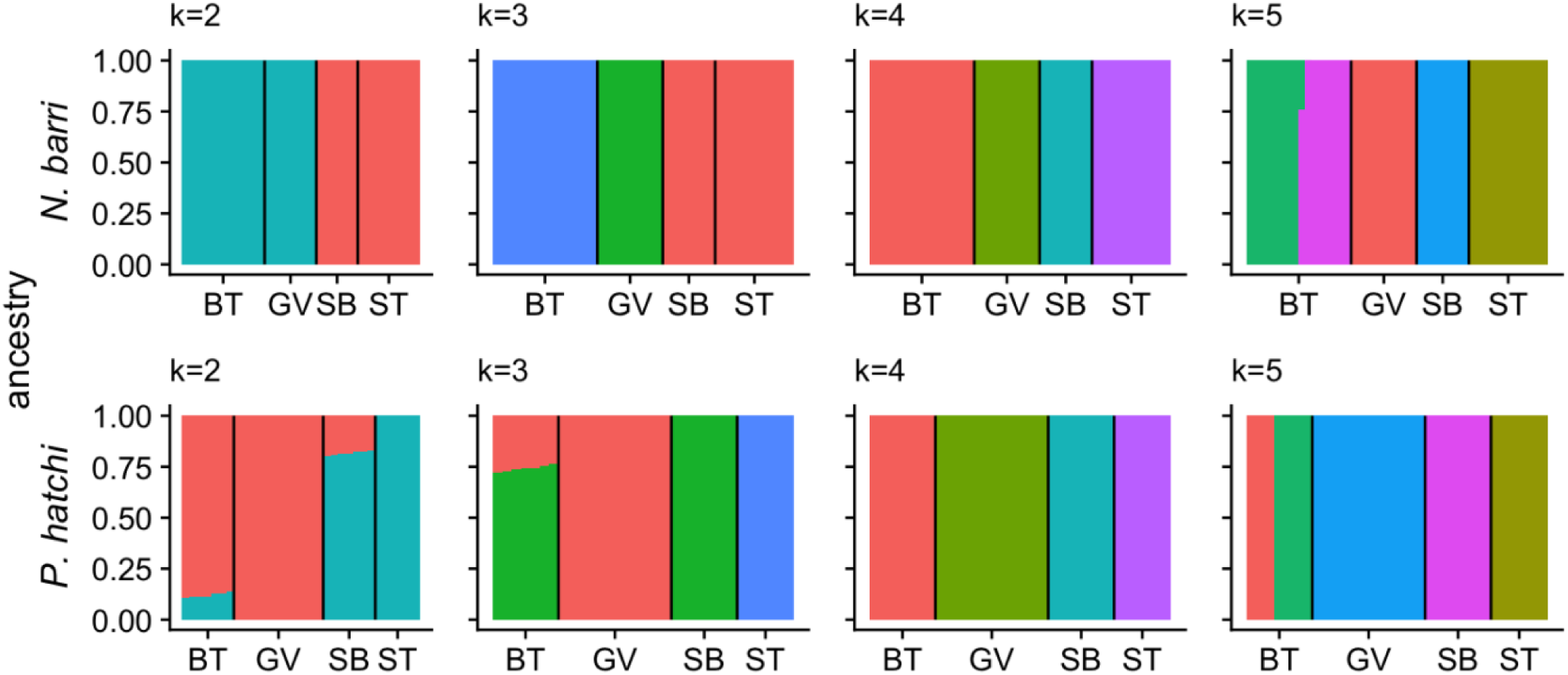
Admixture plots for varying numbers of ancestral populations (k = 2-5) for *N. barri* (top) and *P. hatchi* (bottom). For each panel, k indicates the number of ancestral populations assumed during the analysis and stacked bars represent estimates of proportional ancestry for each individual. Black vertical lines separate groups of individuals from different caves: (BT = Buggytop; GV = Grapevine; SB = Sewanee Blowhole; ST = Solomon’s Temple).

### Genetic differentiation between caves

Genetic differentiation between caves was high. Pairwise weighted F_ST_ (Angsd) ranged from 0.33 to 0.52 for *N. barri*, and 0.3 to 0.36 for *P. hatchi*. Unweighted F_ST_ (Angsd) and Weir and Cockerham’s F_ST_ estimated from hard genotype calls were lower, but still considerable, with a minimum value of 0.12 (Table 2; Fig. 4). Based on hierarchical clustering and admixture analyses (Fig. 2; Fig. 3), we expected to find greater genetic differentiation between caves located in different watersheds. This pattern was consistent for *N. barri*, but not for *P. hatchi* (Fig. 4). Although all the difference estimates were tightly correlated (Fig. S1), only absolute genetic distance (d_XY_ from hard genotype calls) was consistently lower within watersheds for *P. hatchi*.

**Figure 4:**
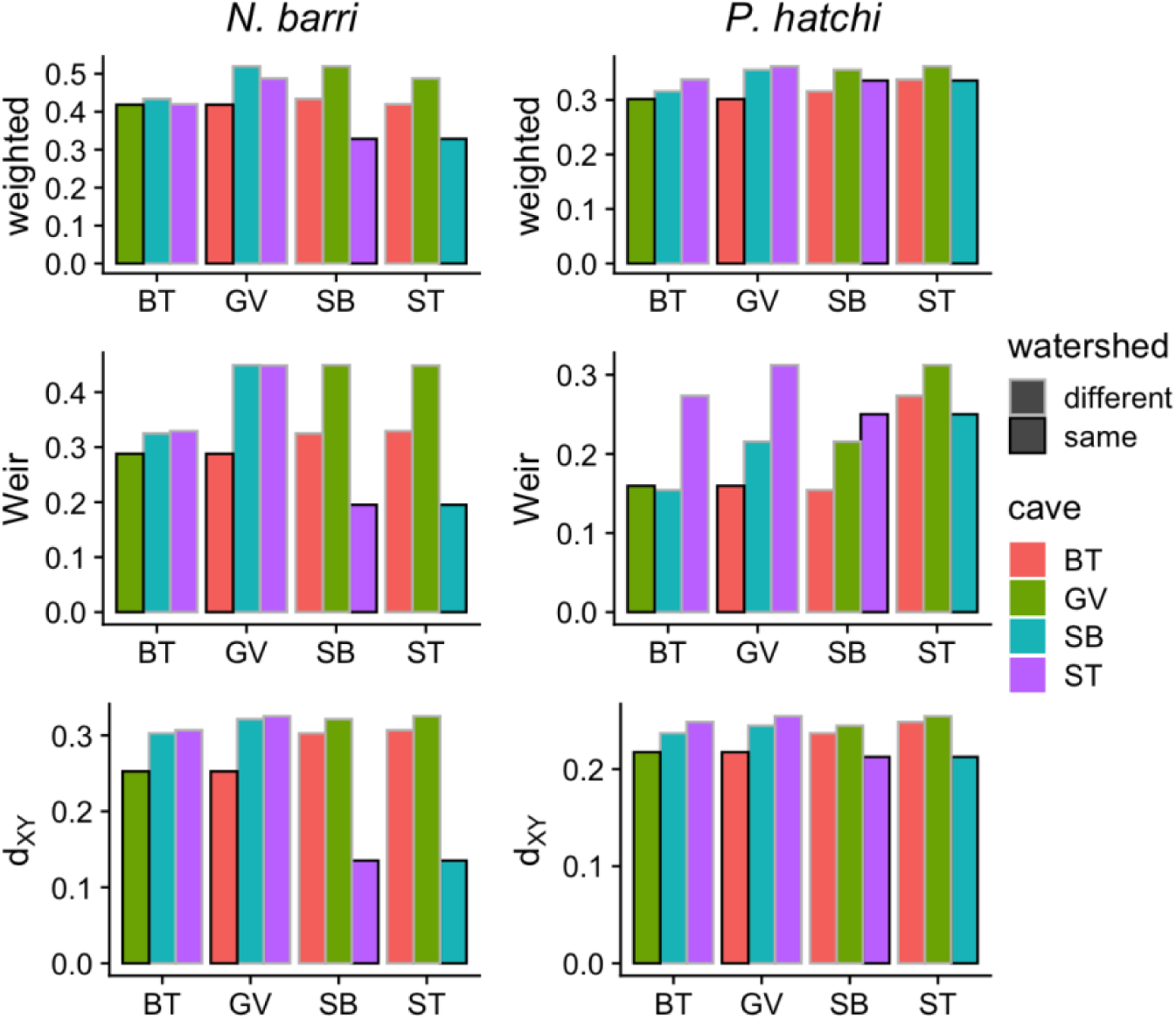
Pairwise estimates of genetic differentiation between caves. The first cave in each pair is indicated on the X axis. The second cave is indicated by the bar color. Whether the two caves are located in the same watershed is indicated by the bar outline color. The statistics are: weighted = weighted F_ST_ computed using Angsd; Weir = Weir and Cockerham’s F_ST_ averaged across all variant sites from hard genotype calls; d_XY_ = absolute genetic distance averaged across all variant sites from hard genotype calls.

### Nucleotide diversity

In both species, estimates of nucleotide diversity based on genotype likelihoods indicated surprising levels of diversity that varied with cave length. For individual caves, per site nucleotide diversity (π) ranged from 1.17e-3 to 2.43e-3. For all but the longest cave (Buggytop Cave; BT), the beetle *P. hatchi* had higher nucleotide diversity than the spider *N. barri* (Table 3). For both species, nucleotide diversity correlated positively with cave length (Fig. 5). Assuming a mutation rate of 2.8e-9 estimated for *Drosophila* (Keightley et al. 2014), the effective population sizes based on the π estimates for *P. hatchi* ranged from 1.4e5 in Solomon’s Temple to 2.2e5 in Buggytop, with a range of 1.0e5 to 2.3e5 for *N. barri* (Table 3). Nucleotide diversity estimates based on hard genotype calls were proportionally similar, but on average 8-fold lower than from genotype likelihoods (Table 3). This likely resulted from greater stringency during filtering of variant sites from the hard genotype calls. Estimates of π from hard genotype calls were similarly positively associated with cave length (Fig. S2).

**Figure 5:**
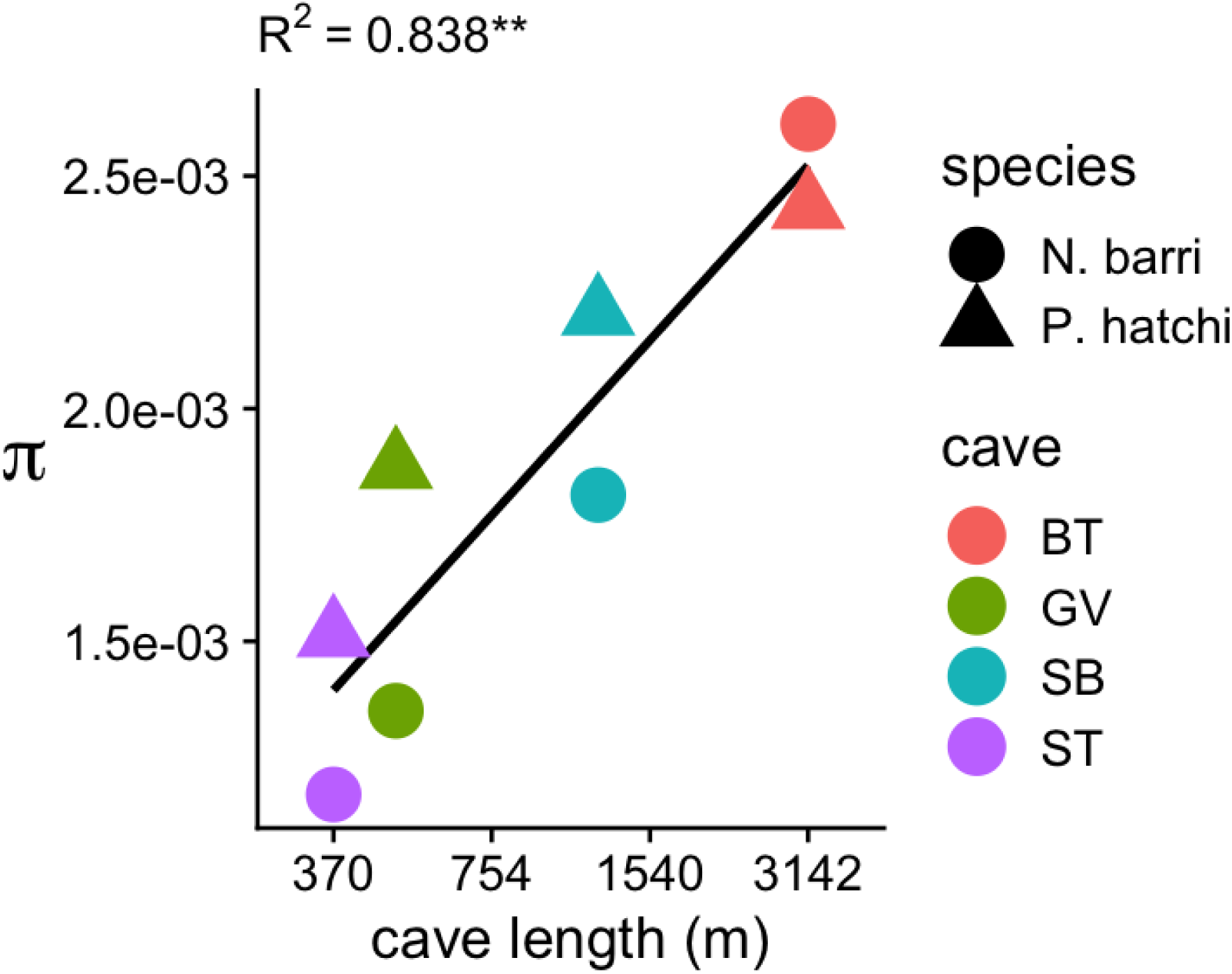
Relationship between per site nucleotide diversity (π) and cave length. The X-axis shows cave length on the log scale. The Y-axis shows the nucleotide diversities for the two species in each cave computed using Angsd. Point color indicates cave and point shape indicates species. Black line traces the linear regression for all points. R^2^ for the linear model for all points is given above the plot (p < 0.01).

**Table 3:** Nucleotide diversity for each species and cave estimated using genotype likelihoods (Angsd) and from hard genotype calls (mpileup).

| Species | Cave | Cave length (m) | $\pi$ (Angsd) | Ne (Angsd) | $\pi$ (mpileup) | Ne (mpileup) |
| --- | --- | --- | --- | --- | --- | --- |
| *N. barri* | ST | 370 | 1.2E-03 | 1.0E+05 | 1.6E-04 | 1.4E+04 |
| *N. barri* | GV | 490 | 1.4E-03 | 1.2E+05 | 2.0E-04 | 1.8E+04 |
| *N. barri* | SB | 1219 | 1.8E-03 | 1.6E+05 | 2.1E-04 | 1.9E+04 |
| *N. barri* | BT | 3142 | 2.6E-03 | 2.3E+05 | 3.3E-04 | 2.9E+04 |
| *P. hatchi* | ST | 370 | 1.5E-03 | 1.4E+05 | 1.2E-04 | 1.0E+04 |
| *P. hatchi* | GV | 490 | 1.9E-03 | 1.7E+05 | 2.7E-04 | 2.4E+04 |
| *P. hatchi* | SB | 1219 | 2.2E-03 | 2.0E+05 | 3.3E-04 | 2.9E+04 |
| *P. hatchi* | BT | 3142 | 2.4E-03 | 2.2E+05 | 3.5E-04 | 3.1E+04 |

## Discussion

We used genome-wide genotyping to examine population structure of two troglobionts from the southern Cumberland Plateau in Tennessee. Despite relatively small distances between caves (no two caves were more than 15 km apart), we detected strong population structure for both species.

Hierarchical clustering, PCA, and Admixture clearly identified each cave as a genetically distinct population (Fig. 2; Fig. 3). Pairwise estimates of genetic differentiation further supported these results, with a minimum weighted F_ST_ of 0.33 for *N. barri* and 0.30 for *P. hatchi* (Table 2), indicating “very great differentiation” (Wright 1978). For comparison, a recent 2bRAD study on the coral *Acropora millepora* across the Great Barrier Reef, including sites located over 1200 km apart, detected a maximum pairwise F_ST_ of 0.014 (Matz et al. 2018). Hence, populations from separate caves are remarkably isolated. These findings corroborate previous population genetic studies using single gene approaches (Snowman et al. 2010; Dixon and Zigler 2011) at the genomic scale.

Under a neutral model, nucleotide diversity is expected to be linearly proportional to the effective population size. For both species, we detected a positive association between nucleotide diversity and cave length (Fig. 5; Fig. S2). While the number of caves in our study was small, this result is consistent with the intuitive idea that larger caves harbor larger, more genetically diverse populations. Indeed, based on the strength of the correlations observed, cave size appears to be a dominant force shaping troglobiont genetic diversity in our study area. Future studies spanning larger numbers of caves will shed light on how generally this pattern occurs.

Based on π estimates, the effective population sizes of both species were surprisingly large (Table 3), especially as caves are expected to harbor relatively small populations (Culver and Pipan 2019). Because Next-Generation sequencing reads originate from single molecules, they are subject to several sources of error such as DNA damage, PCR errors, and sequencing errors, which can inflate estimates of nucleotide diversity (Pool et al. 2010). De novo locus generation likely further contributes to this bias. Fortunately, these error sources can be reasonably expected to influence all samples similarly (Nunziata and Weisrock 2018). Hence, while we believe absolute estimates of Ne reported here are likely inflated, and should be considered cautiously, relative comparisons of diversity are still reliable. This idea is illustrated by the concurrent associations between cave length and π estimated using Angsd and from hard genotypes, despite the roughly 8-fold difference between them in absolute terms (Table 3; Figure 5; Figure S2).

Based on our results, we conclude that gene flow between caves is rare. This is consistent with the inability of troglobionts to traverse even small distances between the caves (Fig. 1). Hence the analogy of caves to islands in a sea of surface habitat holds for these species (Culver and Pipan 2019; Snowman et al. 2010). It is thought that migration of troglobionts must occur via subterranean connections (Culver and Pipan 2019; Trontelj et al. 2019). Consistent with this theory, hierarchical clustering of populations for both species paired caves by watershed, rather than physical distance (Fig. 1; Fig. 2). This possibly reflects greater frequency of rare subterranean connections, or more recent vicariance between caves located in the same watershed. Estimates of genetic differentiation were consistent with this theory for *N. barri*, but not for *P. hatchi*.

Caves provide a unique opportunity for insight into evolutionary processes. For instance, troglomorphic traits have been divided into ‘constructive’ traits, such as appendage and antennae elongation, and ‘regressive’ traits, such as eye and pigment loss, which are hypothesized to vary in both their rates and mechanisms of evolution (Culver et al. 1995). Regressive traits are considered capable of relatively rapid evolution via drift and mutation accumulation, whereas constructive traits are hypothesized to evolve more slowly by positive selection (Culver et al. 1995; Rétaux and Casane 2013). A similar distinction is evident in a passage from Darwin, attributing eye loss to disuse rather than selection because it was “difficult to imagine that the eyes, though useless, could be in any way injurious to animals living in darkness” (Darwin 1959). However, energetic costs have since been cited as a potential selective pressure driving eye loss (Moran et al. 2015), and there is evidence from troglobiotic fish that eye loss involves epigenetic changes, rather than mere failures to develop due to loss-of-function mutations (Gore et al. 2018). These studies, along with the findings reported here strengthen the idea that caves are ideal natural laboratories for evolutionary insight (Poulson and White 1969). The highly restricted migration rates we observe indicate an opportunity to examine ongoing and nearly independent evolution to highly unique conditions. Hence, through further application of genomic tools, these natural laboratories have great potential to inform evolutionary understanding.

## Acknowledgements

This study was supported by the National Science Foundation grant IOS-1755277 to M.V.M. Data analysis was performed with the help of the Texas Advanced Computing Center. We thank Mikhail Matz for support as well as assistance with analysis and composition.

## Data accessibility

Demultiplexed reads for all samples are available on the NCBI SRA database (PRJNA601737). All scripts used for data processing, statistical analysis, and plotting figures, as well as intermediate data files are available on github: https://github.com/grovesdixon/caveRAD.

## References

GitHub (2014) ANGSD. Available from https://github.com/ANGSD/angsd (accessed February 2020)

Buhay JE, Crandall KA (2005) Subterranean phylogeography of freshwater crayfishes shows extensive gene flow and surprisingly large population sizes. Mol Ecol 14:4259–4273

Buhay JE, Moni G, Mann N, Crandall KA (2007) Molecular taxonomy in the dark: Evolutionary history, phylogeography, and diversity of cave crayfish in the subgenus Aviticambarus, genus Cambarus. Mol Phylogenet Evol 42:435–448

Carver LM, Perlaky P, Cressler A, Zigler KS (2016) Reproductive seasonality in Nesticus (Araneae: Nesticidae) cave spiders. PLoS One 11:7–8

Christman MC, Culver DC (2001) The relationship between cave biodiversity and available habitat. J Biogeogr 28:367–380

Christman MC, Culver DC, Madden MK, White D (2005) Patterns of endemism of the eastern North American cave fauna. J Biogeogr 32:1441–1452

Cook LM, Saccheri IJ (2013) The peppered moth and industrial melanism: Evolution of a natural selection case study. Heredity 110:207–212

Culver DC, Kane TC, Fong DW (1995) Adaptation and natural selection in caves: the evolution of Gammarus minus. Harvard University Press, Cambridge, MA

Culver DC, Master LL, Christman MC, Hobbs HH (2000) Obligate Cave Fauna of the 48 Contiguous United States. Conserv Biol 14:386–401

Culver DC, Pipan T (2019) The biology of caves and other subterranean habitats, 2nd edn. Oxford University Press, New York, NY

Danecek P, Auton A, Abecasis G, et al (2011) The variant call format and VCFtools. Bioinformatics 27:2156–8

Darwin C (1959) On The Origin of Species by Means of Natural Selection, or Preservation of Favoured Races in the Struggle for Life. John Murray, London

GitHub (2019) caveRAD. Available from https://github.com/grovesdixon/caveRAD (accessed March 2020).

Dixon G, Kitano J, Kirkpatrick M (2019) The Origin of a New Sex Chromosome by Introgression between Two Stickleback Fishes. Mol Biol Evol 36:28–38

Dixon GB, Davies SW, Aglyamova G V, et al (2015) Genomic determinants of coral heat tolerance across latitudes. Science 348:1460–1462

Dixon GB, Zigler KS (2011) Cave-obligate Biodiversity on the Campus of Sewanee: The University of the South, Franklin County, Tennessee. Northeast Nat 10:251–266

Fumagalli M, Vieira FG, Korneliussen TS, et al (2013) Quantifying population genetic differentiation from next-generation sequencing data. Genetics 195:979–992

Gore A V., Tomins KA, Iben J, et al (2018) An epigenetic mechanism for cavefish eye degeneration. Nat Ecol Evol 2:1155–1160

Guo Y, Li J, Li CI, et al (2012) The effect of strand bias in Illumina short-read sequencing data. BMC Genomics 13:1–11

Hahn MW (2018) Molecular Population Genetics, 1st edn. Oxford University Press, New York, NY

Hedin M, Dellinger B (2005) Descriptions of a new species and previously unknown males of Nesticus (Araneae: Nesticidae) from caves in Eastern North America, with comments on species rarity. Zootaxa 19:1–19

Hedin MC (1997) Speciational history in a diverse clade of habitat-specialized spiders (Araneae: Nesticidae: Nesticus): Inferences from geographic-based sampling. Evolution 51:1929–1945

Keightley PD, Ness RW, Halligan DL, Haddrill PR (2014) Estimation of the spontaneous mutation rate per nucleotide site in a Drosophila melanogaster full-sib family. Genetics 196:313–320

Korneliussen TS (2013) ANGSD. Available from http://www.popgen.dk/angsd/index.php/ANGSD (accessed February 2020)

Korneliussen TS, Albrechtsen A, Nielsen R (2014) ANGSD: Analysis of Next Generation Sequencing Data. BMC Bioinformatics 15:1–13

Korneliussen TS, Moltke I, Albrechtsen A, Nielsen R (2013) Calculation of Tajima’s D and other neutrality test statistics from low depth next-generation sequencing data. BMC Bioinformatics 14

Langmead B, Salzberg SL (2012) Fast gapped-read alignment with Bowtie 2. Nat Methods 9:357–9

Leray VL, Caravas J, Friedrich M, Zigler KS (2019) Mitochondrial sequence data indicate “Vicariance by Erosion” as a mechanism of species diversification in North American Ptomaphagus (Coleoptera, Leiodidae, Cholevinae) cave beetles. 57:35–57

Li H (2011) A statistical framework for SNP calling, mutation discovery, association mapping and population genetical parameter estimation from sequencing data. Bioinformatics 27:2987–2993

Li H, Handsaker B, Wysoker A, et al (2009) The Sequence Alignment/Map format and SAMtools. Bioinformatics 25:2078–2079

Li W, Godzik A (2006) Cd-hit: A fast program for clustering and comparing large sets of protein or nucleotide sequences. Bioinformatics 22:1658–1659

Matz M V., Treml EA, Aglyamova G V., Bay LK (2018) Potential and limits for rapid genetic adaptation to warming in a Great Barrier Reef coral. PLoS Genet 14:1–19

GitHub (2019) 2bRAD_denovo. Available from https://github.com/z0on/2bRAD_denovo (accessed February 2020).

Moran D, Softley R, Warrant EJ (2015) The energetic cost of vision and the evolution of eyeless Mexican cavefish. Sci Adv 1

Nei M, Li W-H (1979) Mathematical model for studying genetic variation in terms of restriction endonucleases. Proc Natl Acad Sci USA 76:5269–5273

Nielsen R, Korneliussen T, Albrechtsen A, et al (2012) SNP calling, genotype calling, and sample allele frequency estimation from new-generation sequencing data. PLoS One 7

Niemiller ML, Zigler KS (2013) Patterns of Cave Biodiversity and Endemism in the Appalachians and Interior Plateau of Tennessee, USA. PLoS One 8

Nunziata SO, Weisrock DW (2018) Estimation of contemporary effective population size and population declines using RAD sequence data. Heredity 120:196–207

Peck SB (1986) Evolution of adult morphology and life-history characters in cavernicolous Ptomaphagus beetles. Evolution 40:1021–1030

Polo-Cavia N, Gomez-Mestre I (2017) Pigmentation plasticity enhances crypsis in larval newts: Associated metabolic cost and background choice behaviour. Sci Rep 7:1–10

Pool JE, Hellmann I, Jensen JD, Nielsen R (2010) Population genetic inference from genomic sequence variation. Genome Res 20:291–300

Porter ML (2007) Subterranean biogeography: What have we learned from molecular techniques? J Cave Karst Stud 69:179–186

Poulson TL, White WB (1969) The Cave Environment. Science 165:971–981

Rétaux S, Casane D (2013) Evolution of eye development in the darkness of caves: Adaptation, drift, or both? Evodevo 4:1–12

Reynolds J, Weir BS, Cockerham CC (1983) Estimation o f the coancestry coefficient: basis for a short-term genetic distance. Genet Soc Am 105:767–779

Rokas A, Abbot P (2009) Harnessing genomics for evolutionary insights. Trends Ecol Evol 24:192–200

Skotte L, Korneliussen TS, Albrechtsen A (2013) Estimating individual admixture proportions from next generation sequencing data. Genetics 195:693–702

Snowman C V., Zigler KS, Hedin M (2010) Caves as islands: mitochondrial phylogeography of the cave-obligate spider species Nesticus barri (Araneae: Nesticidae). J Arachnol 38:49–56

Trontelj P, Borko Š, Delić T (2019) Testing the uniqueness of deep terrestrial life. Sci Rep 9:1–9

Vieira FG, Fumagalli M, Albrechtsen A, Nielsen R (2013) Estimating inbreeding coefficients from NGS data: Impact on genotype calling and allele frequency estimation. Genome Res 23:1852–1861

Wang S, Meyer E, McKay JK, Matz M V (2012) 2b-RAD: a simple and flexible method for genome-wide genotyping. Nat Methods 9:808–810

Weir BS, Cockerham CC (1984) Estimating F-Statistics for the Analysis of Population Structure. Evolution 38:1358–1370

Wright S (1978) Variability Within and Among Natural Populations. University of Chicago Press, Chicago

Zigler KS, Niemiller ML, Fenolio DB (2014) 2014 NSS Convention Guidebook: Alabama. Huntsville, Alabama

