## Supplemental Figures for "Testing the ‘caves as islands’ model in two cave-obligate invertebrates with a genomic approach"

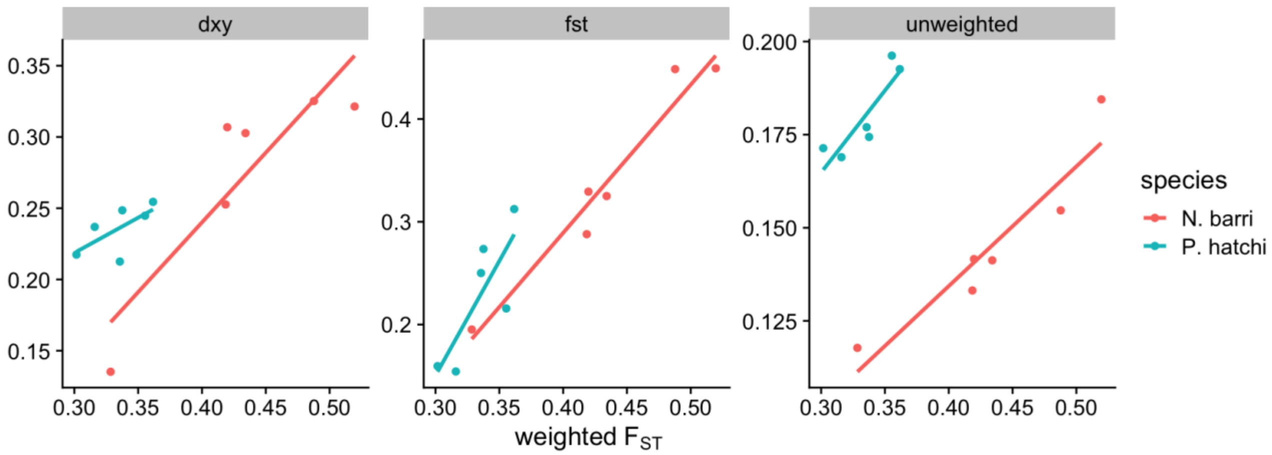


Figure S1: Correlation between pairwise estimates of genetic differentiation. The X axes show weighted $F_{ST}$ computed using Angsd. The Y axes show the statistic indicated in the panel title: dxy = absolute genetic distance averaged across all variant sites from hard genotype calls; fst = Weir and Cockerham's $F_{ST}$ averaged across all variant sites from hard genotype calls; unweighted = unweighted $F_{ST}$ computed using Angsd.

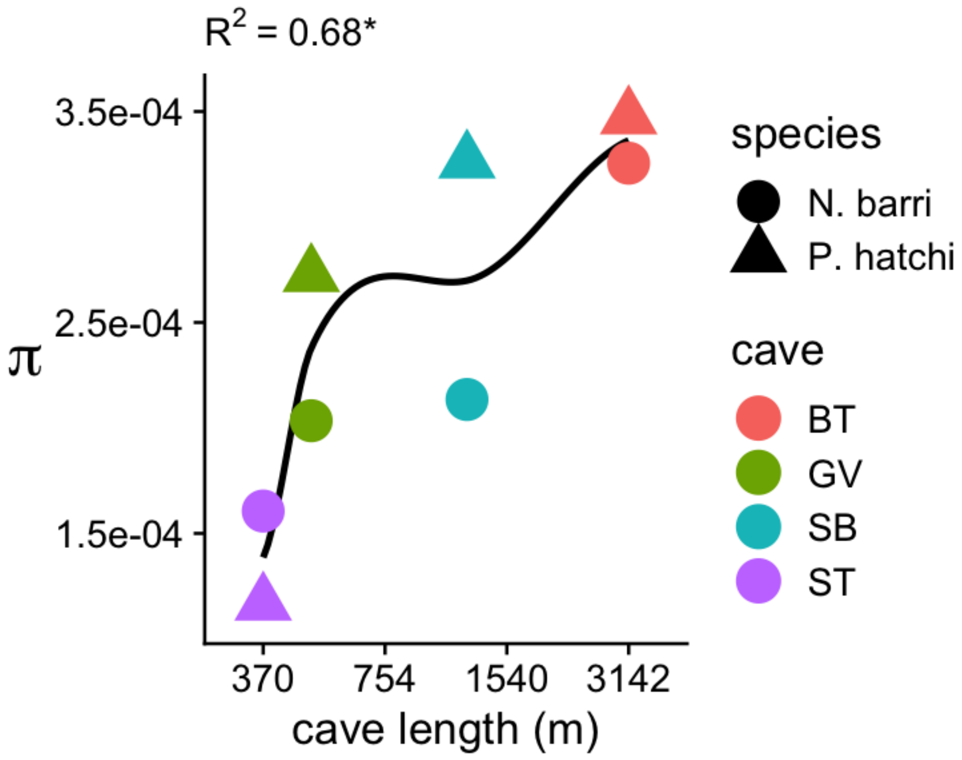


Figure S2: Relationship between per site nucleotide diversity (π) and cave length. X axis cave length on the log scale. Y axis shows the nucleotide diversities for each species caves pair computed from hard genotype calls. Point color indicates cave and point shape indicates species. Black line traces a locally smoothed regression for all points. $R^2$ for the linear model for all points is given above the plot ($p < 0.01$).
